# Convergent Multimodal Evidence of Cortical Excitation-Inhibition Imbalance in Psychosis

**DOI:** 10.64898/2026.03.31.715583

**Authors:** Ioana Varvari, Max Doody, Zilin Li, Dominic Oliver, Philip McGuire, Matthew M. Nour, Robert A. McCutcheon

## Abstract

Psychosis is increasingly understood as a disorder of disrupted cortical excitation-inhibition balance, yet robust non-invasive translational biomarkers remain lacking. The resting-state fMRI Hurst exponent (HE) and EEG aperiodic spectral exponent are promising complementary biomarkers, with lower values in each proposed to reflect a shift towards cortical hyperexcitability, but they have not been jointly examined in psychosis, and the spatial and molecular architecture of HE alterations remains poorly defined. We therefore tested for convergent systems-level signatures across independent cohorts and modalities, using resting-state fMRI (107 patients, 53 controls) and EEG (547 patients, 363 controls). Whole-brain and regional HE were estimated using wavelet methods, and EEG aperiodic exponents were quantified using spectral parameterisation. Compared with healthy controls, individuals with psychosis showed reduced whole-brain HE and widespread regional reductions. Regional HE case-control differences were associated with cortical gene-expression patterns, with enrichment for potassium channel and GABA receptor pathways, and correlated with noradrenergic, muscarinic, serotonergic, glutamatergic and dopaminergic receptor density maps, but not with cortical thickness or symptom or cognitive measures. In the independent EEG cohort, psychosis was similarly associated with a reduced aperiodic spectral exponent. Together, these findings provide cross-modal evidence for altered cortical resting-state dynamics in psychosis, consistent with a shift towards cortical hyperexcitability. Integration with receptor-density and transcriptomic maps implicates biologically plausible molecular pathways and supports HE and EEG aperiodic activity as scalable translational biomarkers in psychosis.

## Introduction

Excitation - inhibition (E/I) balance is a fundamental property of neuronal circuits and is essential for normal brain function. Excitatory (glutamatergic) and inhibitory (GABAergic) inputs dynamically interact to regulate neuronal firing and network activity across brain states. GABAergic inhibitory interneurons play a central role in maintaining this balance and prevent pathological extremes such as hyperexcitable or silent brain states (1). Disruption of E/I balance has been proposed as a core pathophysiological mechanism underlying psychosis, potentially accounting for features observed at a range of temporal and spatial scales. Perturbations of E/I balance can alter mesoscopic neuronal oscillations detectable using EEG, which in turn may contribute to large-scale network dysconnectivity that is measurable with fMRI (2–6). These disturbances may manifest clinically as positive (3), cognitive (2, 4, 7), and negative (8) symptoms of psychosis, which contribute to the functional and social impairments associated with the disorder (9).

Despite broad recognition of E/I imbalance, most evidence linking microcircuit dysfunction to large network disruption is largely derived from ex vivo, in silico, and in vivo animal studies (2–4, 10). Direct measurement of E/I shifts in humans remaining challenging, and translational evidence in patients with psychosis is comparatively limited. Traditional noninvasive neuroimaging metrics such as EEG power spectra, functional magnetic resonance imaging fMRI, magnetic resonance spectroscopy (MRS) metabolites, provide indirect, relatively nonspecific proxies with limited validation against microcircuit mechanisms (10–12). This gap highlights the need for noninvasive translational biomarkers.

Two emerging candidates are the Hurst Exponent (HE) for rs-fMRI (13–16) and the aperiodic 1/f spectrum for the EEG (17), both predicted by computational microcircuit models and empirically linked to changes in E/I shifts across animal and human studies. The HE asks, *“Does current brain activity depend on its past?’’* (13), quantifying the persistence of temporal dependencies, with lower values indicating reduced inhibition and increased cortical excitability (14, 18). The aperiodic 1/f slope captures the broadband spectral structure of neural activity, with lower values similarly indicating a shift toward reduced inhibitory drive (19). Together, these different metrics capture E/I shifts from complimentary perspectives, offering a translational framework for psychosis research.

To date, these metrics have not been comprehensively characterised in psychosis. In this study, we investigated the HE and the aperiodic 1/f spectrum as proxy measures of E/I balance in two independent clinical psychosis datasets. We estimated HE using robust wavelet-based methods to capture long-range temporal dependencies, and quantified the aperiodic 1/f slope (14). Our aims were to test case-control differences in these measures, characterize their spatial patterns, and examine their associations with clinical features and putative biological substrates.

## Materials and methods

### Population

fMRI data were obtained from the Human Connectome Project for Early Psychosis (HCP-EP) release 1.1 and EEG data were obtained from the Bipolar Schizophrenia Network on Intermediate Phenotypes-2 (BSNIP2). The HCPEP dataset provided data from 183 patients with psychosis within three years from illness onset and 68 demographically matched healthy controls, diagnosed with the Structured Clinical Interview for DSM-5 (SCID-5-RV). The BSNIP2 dataset provided data from 582 patients with a diagnosis of longstanding (more than 5 years from 1^st^ admission) schizophrenia or schizoaffective disorder and 379 healthy controls, diagnosed with the Structured Clinical Interview for DSM-IV (SCID-IV). Detailed inclusion and exclusion criteria are described elsewhere (20, 21). All participants provided written informed consent under institutional review board approved procedures.

### Data collection

HCP-EP fMRI data were acquired on a 3T Siemens Connectome Skyra scanner using multiband echo - planar imaging (TR = 0.8 s, TE = 33 ms, flip angle = 52°, voxel = 2 mm, 60 slices, multiband factor = 8). Participants completed four 5 min eye open runs. Psychopathology was assessed with the Positive and Negative Syndrome Scale (PANSS) and cognitive performance was measured using the NIH Toolbox Cognitive Battery. BSNIP2 EEG data were recorded using the Compumedics Neuroscan 64-channel Quik-Cap with a sampling rate 1000 Hz and no online band-pass filtering. Psychopathology was assessed with PANSS, and cognitive functioning with the Brief Assessment of Cognition in Schizophrenia (BACS) battery. Detailed data collection protocols are described elsewhere (20, 21).

### Data preprocessing

fMRI data were preprocessed with fMRIPrep v25.0 (22), then denoised using wavelet despiking (BrainWavelet toolbox v2.0, threshold = 10) (23) and 13 parameter nuisance regression (6 motion parameters, their first derivatives, and cerebrospinal fluid signal) following prior work validating HE estimation (14). Rs - time series were parcellated into 360 cortical and 66 subcortical regions, using the HCPex atlas v1.1. (24) EEG data were preprocessed with an MNE based pipeline including average re-referencing, notch and 0.5 - 40 Hz band-pass filtering, bad-channel interpolation, Independent Component Analysis (ICA) based ocular artefacts removal and 2 second epochs segmentation (25). Power spectral densities (PSDs) were averaged per participant to yield subject-level spectra. Aperiodic 1/f components were fit using Spectral Parameterization (FOOOF) across 1 - 40 Hz (26). See eSection 1.1 for detailed preprocessing details.

### Data analysis

Rs-fMRI analyses were implemented in Python v 3.12. Regional HE was computed with a wavelet-based maximum likelihood estimator (14). A whole-brain HE was derived by averaging regional HEs. To minimise site effects, regional HE and whole brain HE were harmonized using ComBat, with age, sex, and diagnosis as covariates of interest (27). We implemented automated quality control assessments for quantity of missing data and outliers and excluded participants with mean Framewise Displacement (mFD) > 0.25 mm (see supplement for details).

Group differences in whole brain and regional HE were assessed using ordinary least squares regression (OLS) adjusted for covariates (HE = *β*□ + *β*□Phenotype + *β*□Age + *β*□Sex + *β*□FD + ε). Additionally, phenotype-by-age and phenotype-by-sex interaction models tested moderation effects. Group difference effect sizes were quantified using Cohen’s d. The potential effect of antipsychotic exposure was assessed by comparing whole-brain HE between medicated and unmedicated patients using an independent-samples t-test, and by calculating the Pearson correlation between chlorpromazine-equivalent dose and whole-brain HE. To relate regional HE to PANSS symptoms and cognitive performance, we used 10 fold cross validation (CV) partial least squares regression (PLSR) models across multiple predictor-outcome configurations (Supplementary Table 1). Cognitive performance was summarised using a composite score derived from the NIH Toolbox and BACS batteries.

We then investigated the nature of the regional case-control differences. Parcel-wise effects were examined for enrichment within canonical Yeo 7 networks, to contextualise spatial patterns (28). To explore potential biological correlates, regional HE was Pearson correlated across subjects with similarly but independently harmonised cortical thickness (CT). The spatial distribution of regional HE effects were then related to gene expression profiles from the Allen Human Brain Atlas (29, 30) employing 10 fold CV PLSR, after preparing the gene expression data via the abagen pipeline (31), followed by ontology enrichment with GOrilla (32). Association of the regional HE effects with receptor density distributions were also examined using PLSR and the Hensen Atlas (33). In these analyses, parcel-level CT, gene expression, or receptor density were used as predictors to estimate spatial pattern of the case-control difference in HE. Model interpretation and feature contribution were assessed using variable importance in projection (VIP) scores. See eSection 2 for methodological details.

EEG analyses were implemented in Python (v3.6.8). Following quality control assessments outlier or poor-fit spectra were excluded (see eSection 2). For each participant, all valid sessions were included. Preprocessing and PSD derivation were performed separately for each run, after which a single mean 1/f per participant was calculated. Group-level differences in 1/f aperiodic exponent were assessed using OLS regression controlling for age, sex, offset and site. The 1/f offset is less mechanistically understood than the HE, but can be interpreted as the overall broadband (baseline) level of neuronal activity. Offset effects were similarly tested i.e. controlling for age, sex, 1/f, and site.

Throughout, significance was assessed using Benjamini Hochberg False Discovery Rate (FDR) correction (q < .05 two tailed) and case-control permutation testing (*n* = 10,000). Detailed QC and analysis details are provided in eSections 1.2-1.3.

## Results

### Demographic characteristics

Of 183 participants in the HCP-EP rs-fMRI cohort, 23 were excluded (8 for unavailable imaging, 14 for excessive motion [mFD > 0.25], and 1 outlier > 3 SD), leaving 160 for analysis. Including the outlier showed unchanged results (Supplementary Table 3). In the BSNIP2 EEG cohort 63 of 961 participants were excluded (44 poor spectral fits [R^2^ < 0.8 or exponent/offset out of range], 7 missing age/site data). Demographics are summarised in Table 1.

**Table 1.**
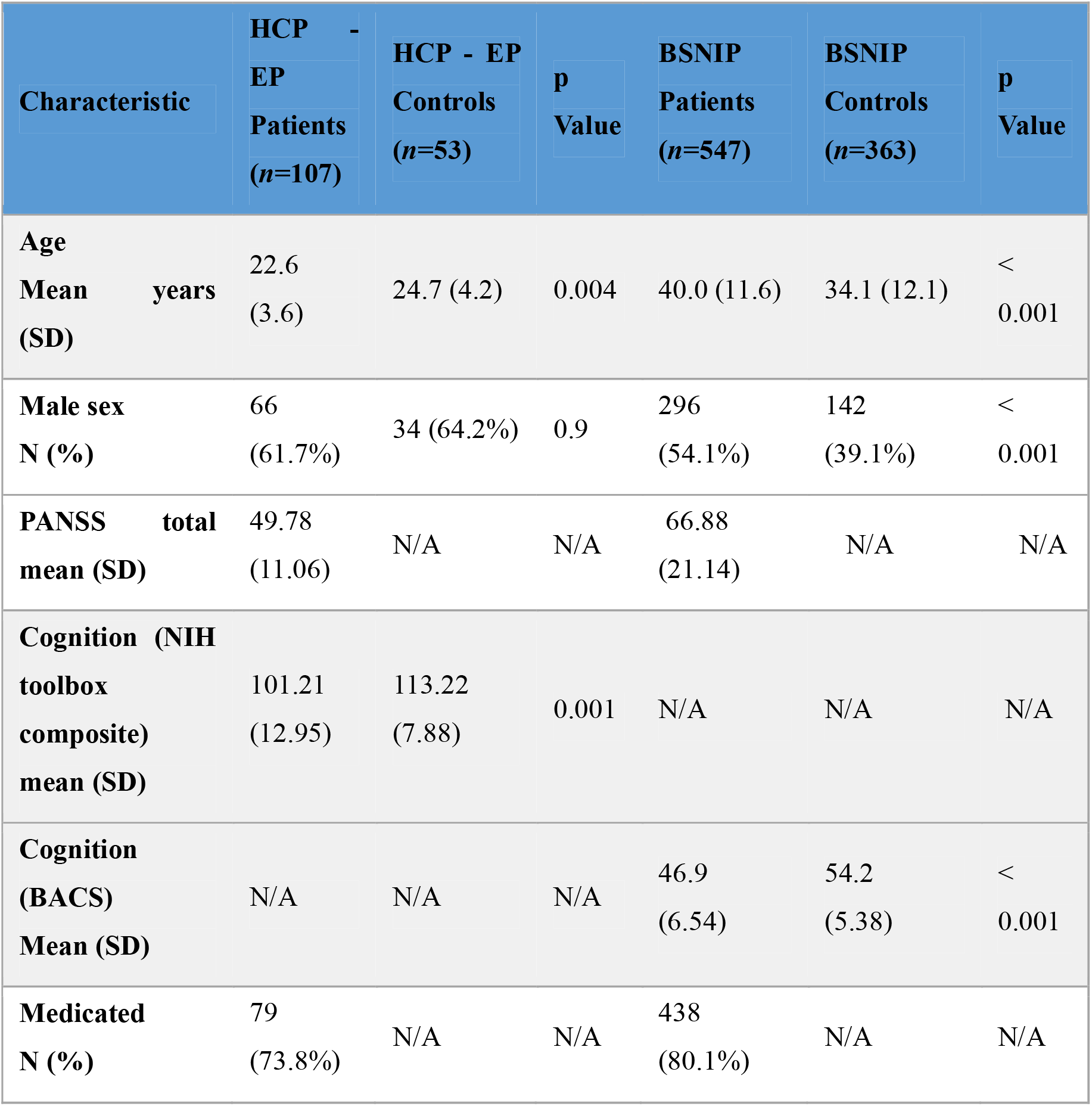
Legend: SD = standard deviation, no = number, PANSS = Positive and Negative Symptoms Scale, N/A= not applicable, cognition total = standardised scored of the total NIH toolbox cognitive battery in HCPEP and BACS battery in BSNIP2. The p values were calculated using independent-samples t-tests for continuous variables (age, cognition) and chi-square tests for categorical variables (sex, ethnicity). For ethnicity, the p values were calculated between proportion of white and ethnical minorities between cases and controls.

### rs-fMRI analyses

#### Whole brain and regional Hurst Exponent group differences

Early psychosis participants showed lower whole brain mean rs-fMRI HE values than controls. Group differences remained significant after adjusting for age, sex, and mean FD (*β* = −0.031; 95% CI = [−0.048, −0.015]; Cohen’s *d* = −0.68, perm *P* < 0.001) (Fig. 1A). HE did not differ significantly between patients taking antipsychotic medications and those who were medication free (p=.66), and did not correlate with chlorpromazine equivalent dose (*r*_*ρ*_ = −0.15, *P* = 0.2) (Fig. 1B-C). No diagnosis × age (*β* = −0.0000; *P* = 0.89) or diagnosis × sex (*β* = 0.0016; *P* = 0.92) interactions were found.

**Figure 1.**
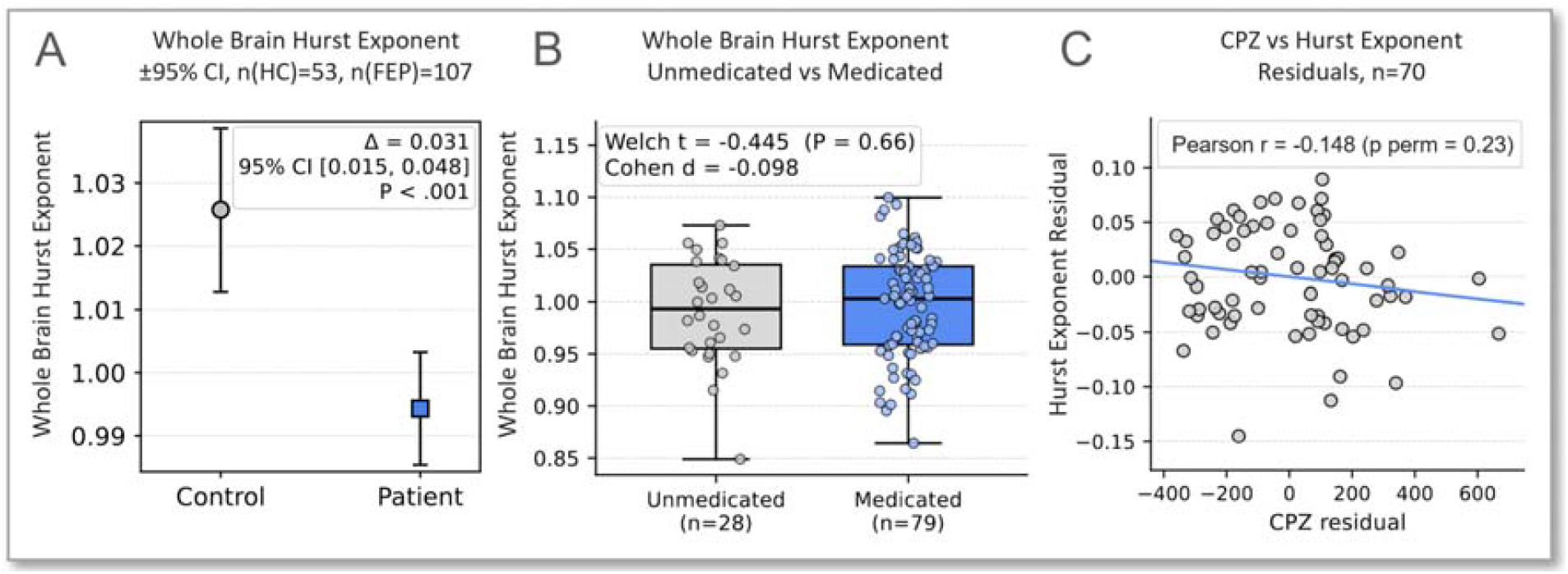
Whole Brain Group Differences and Medication effects on HE. **(A)** Whole Brain HE (±95% CI) for healthy controls (HC; *n* = 53) and first-episode psychosis patients (FEP; *n* = 107). **(B)** Whole Brain HE in unmedicated (*n* = 28) and medicated (*n* = 79) patients, shown as box plots with median and interquartile range. **(C)** Association between chlorpromazine (CPZ) equivalent dose residuals and Whole Brain HE residuals among medicated patients (*n* = 70). Pearson correlations are displayed, with a fitted linear regression line. Hurst residuals represent individual Whole Brain HE values adjusted for covariates (age, sex, and head motion) using linear regression.

There were also widespread cortical HE reductions, at the regional level in early psychosis (223/360 [62%] regions nominally significant, 117 [33%] FDR corrected), which were strongest in somatomotor, insular-opercular, midcingulate, dorsolateral prefrontal, and superior parietal regions (Fig. 2A). There were additional reductions in, 29/66 [44%] regions (22 [33%] FDR corrected), particularly in the thalamus bilaterally (Fig. 2B). The 20 most significant cortical and subcortical regions are listed in Supplementary Table 2.

**Figure 2.**
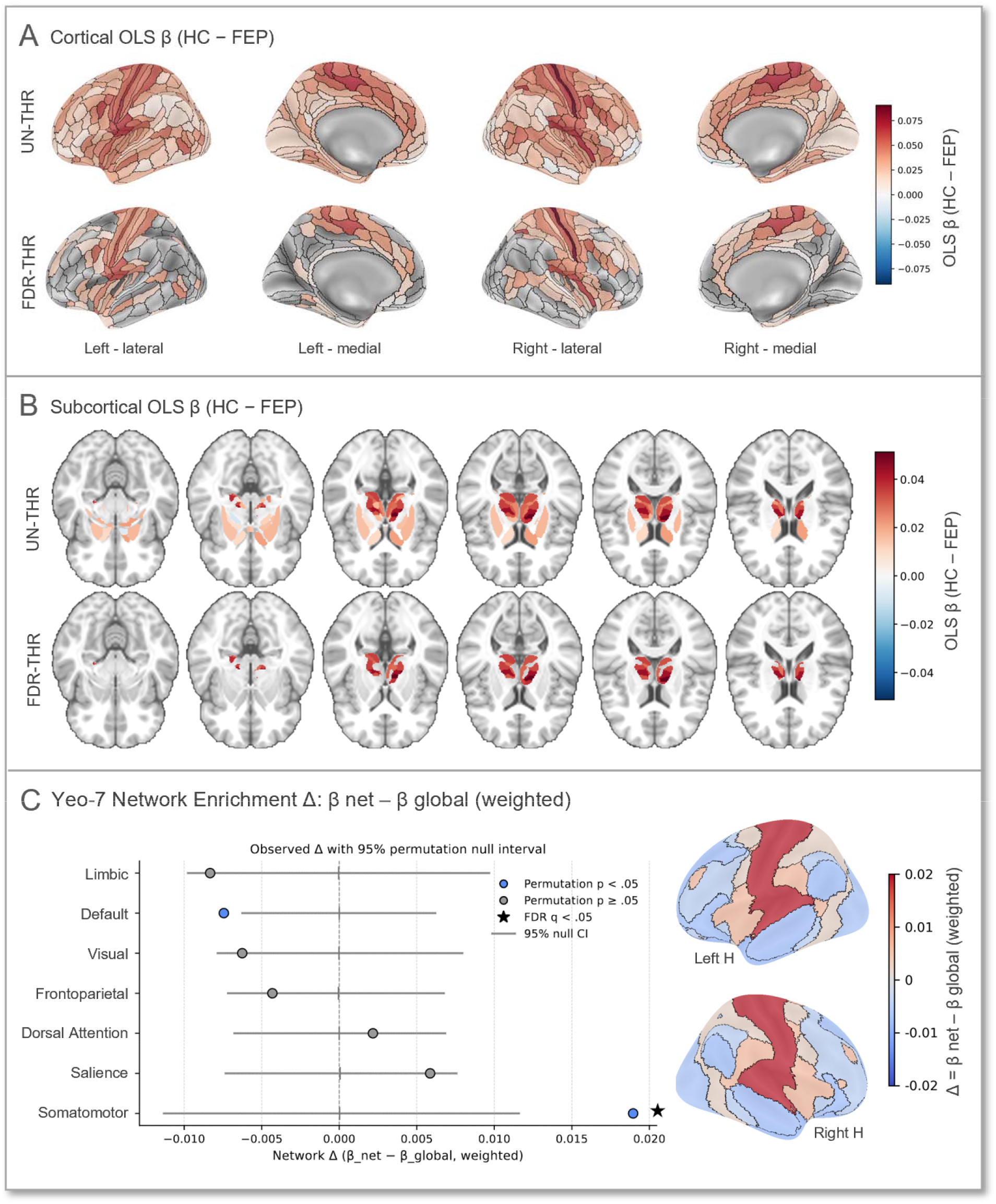
Spatial Nature of Illness Effects. **(A)** Cortical OLS *β* maps showing group differences in regional Hurst exponent between healthy controls (HC) and first-episode psychosis patients (FEP). **(B)** Subcortical OLS *β* maps for the same contrast, displayed on coronal and axial slices. For A and B, warm colours indicate regions where the HE is lower in FEP than HC, reflecting increased excitation. **(C)** Yeo −7 network enrichment analysis showing the weighted difference between network-specific *β* values and the global mean. Points represent observed effects with 95% confidence intervals from 10.000 permutations; filled blue circles indicate networks with permutation *P* < 0.05. *β* values represent OLS effect estimates for HC - FEP contrasts. Maps are thresholded using the Benjamini - Hochberg false discovery rate (FDR) correction at *q* < 0.05, which was applied on permutation p results.

#### Clinical correlates

Across all 10 fold CV PLSR models, there were no statistically significant associations between HE values and symptoms or cognitive scores (all *P* > 0.05).

#### Network enrichment via Yeo 7 functional network atlas

Psychosis related HE reductions were not uniformly distributed across resting state networks. The Somatomotor Network showed the strongest enrichment (Δ*β* = 0.019; perm *P* = 0.001, FDR *q* < 0.05). While not statistically significant, overlap with the salience network was also numerically overrepresented (Δ*β* = 0.006; perm *P =* 0.12), while there was an underenrichment of the default mode network (Δ*β* = −0.005; perm *P* = 0.02, FDR *q* > 0.05) (Fig. 2C).

#### Exploratory analyses of regional HE and biological substrates

We then examined the relationship between HE and cortical thickness. After additional structural MRI QC, 145/160 participants were included (13 excluded for missing CT; 2 excluded as outliers as values >3SD from group mean). Across 360 cortical parcels, HE showed widespread reduction, whereas CT reductions were more limited (18 significant; 9 FDR corrected). We examined correlations across individuals per parcel, and across parcels. No significant correlation between HE and CT across individuals after FDR correction (*q* > 0.05). Adjusted for covariates, regional HE and CT effects, correlated modestly across cortical parcels but did not reach statistical significance when accounting for spatial autocorrelation via permutation testing (*r*_*ρ*_ =0.149, perm *P* = 0.13) (Supplementary Fig. 3). Sensitivity analysis including outliers was unchanged (Supplementary Table 3).

To investigate the relationship between HE and putative neuromodulator and neurotransmitter pathways, we compared the HE spatial distribution to monoaminergic, cholinergic, glutamatergic, GABAergic, and endocannabinoid targets from 19 receptor density maps (33). A 10 fold CV PLSR model showed that receptor distribution patterns significantly predicted regional HE effects. This was statistically significant when assessed against 10,000 random permutations of case-control status (*r*_*ρ*_ = 0.69, perm *P* = 0.01; Fig. 3A). We next examined the model, with bootstrap (B = 10,000) derived VIP scores identifying NET, VAChT, 5-HT□, 5-HT□A, NMDAR, and D□ (VIP > 1) as the strongest contributors (Fig. 3B).

**Figure 3.**
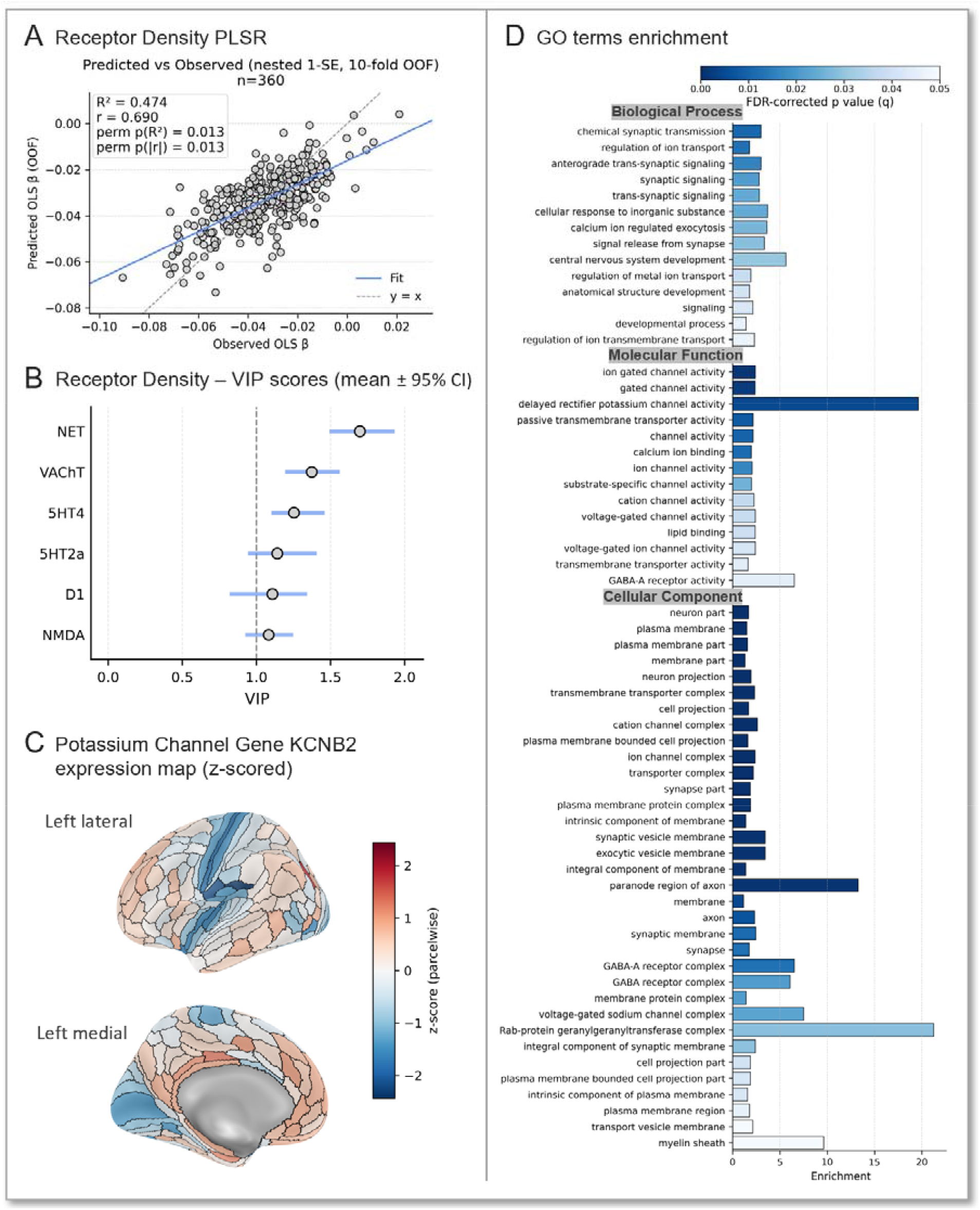
Biological Substrates of Illness Effects. **(A)** Observed versus predicted HE (*β*) from the primary PLSR model based on receptor expression; each point represents out-of-fold predictions from 10-fold cross-validation (*n* = 360). The solid line shows the best-fit regression, and the dashed line indicates the identity line (y = x). **(B)** Receptors with mean VIP values > 1 were considered to contribute above average to the model. Confidence intervals (CI) illustrate receptor importance stability estimates across 10,000 bootstrap resampling; **(C)** KCNB2 expression (left cortex). AHBA expression mapped to HCPex and z-scored across parcels; red = relatively higher, blue = relatively lower, 0 = parcel-mean expression. **(D)** Gene sets surviving FDR corrected (*q* < 0.05) gene enrichment analyses. Bars show enrichment scores across gene categories, with colour indicating FDR corrected significance.

To investigate the relationship between HE and gene expression maps, we compared the HE spatial distribution with regional gene expression profiles derived from the Allen Human Brain Atlas. Using a 10 folds CV PLSR model, we found that cortical gene expression patterns predicted cortical HE patterns. This was statistically significant when tested against 10,000 random permutations of case-control status (*r*_*ρ*_ = 0.55, perm *P* = 0.02) (Supplementary Fig. 5). We next examined the model, with bootstrap (B = 10,000) derived VIP scores showing FDR corrected enrichment (*q* < 0.05) for synaptic signaling, ion transport, and central nervous system development. Enriched cellular components included GABA receptors and voltage-gated sodium channels, while molecular functions were dominated by potassium and calcium channel activity, transmembrane transporter and actin biding (Fig. 3D). Potassium-channel genes showed the strongest enrichment, with KCNB2 exhibiting the strongest spatial correspondence with HE maps (Fig. 3C), indicating that regions with lower KCNB2 expression exhibited larger illness-related differences (*r*_*ρ*_ = −0.46, perm *P* = 0.004; Supplementary Fig. 7).

### EEG findings

Individuals with chronic psychosis showed a lower 1/f exponent than healthy controls, after adjustment for age, sex, site, and offset (*β* = −0.101; 95% CI = [−0.160, −0.042]; Cohen’s *d* = −0.22; perm *P* < 0.001; Fig. 4). The reduced spectral exponent in psychosis indicates flatter aperiodic slopes, consistent with a shift in E/I balance toward excitation. No significant effects of age, sex, or medication were detected. We also examined the aperiodic 1/f offset, which was significantly lower in healthy controls than in chronic psychosis (*β* = −0.179; 95% CI = [−0.303, −0.054]; perm *P* = 0.005; Cohen’s *d* = −0.18; Supplementary Fig. 8). PLSR models for symptoms and cognition found similar results as with rs-fMRI (all *P* > 0.05).

**Figure 4.**
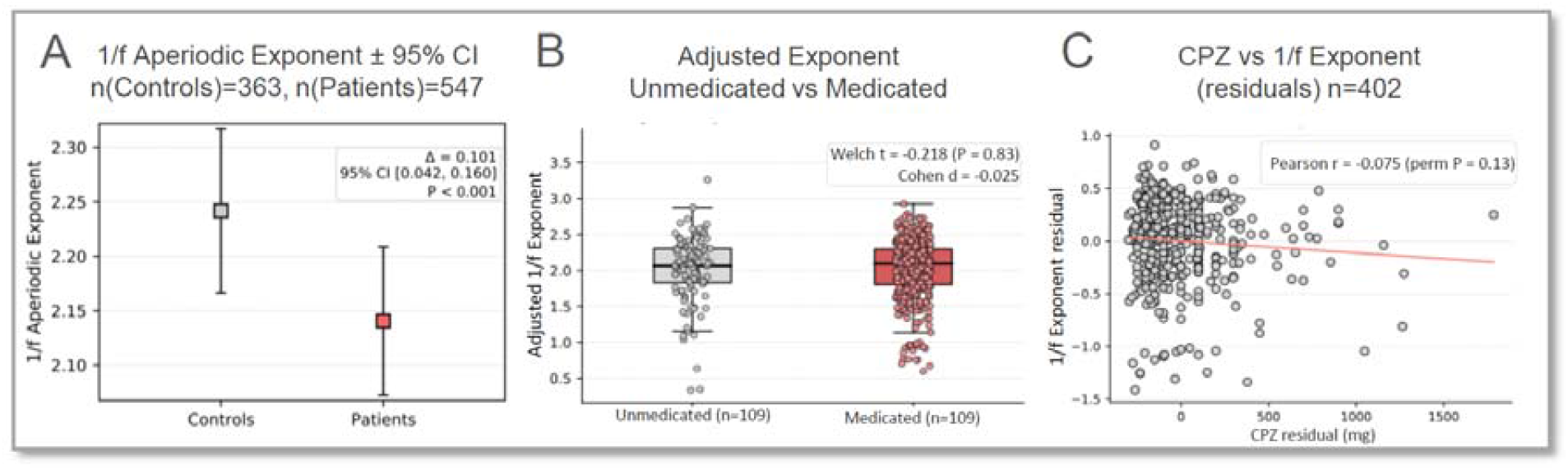
Whole Brain Group Differences and Medication effects on 1/f Aperiodic Exponent. **(A)** 1/f Aperiodic Exponent (±95% CI) for healthy controls (HC; *n* = 363) and schizophrenia patients (SZ; *n* = 547). **(B)** 1/f Aperiodic Exponent in unmedicated (*n* = 109) and medicated (*n* = 438) patients, shown as box plots with median and interquartile range. C) Association between chlorpromazine (CPZ) equivalent dose residuals and 1/f Aperiodic Exponent residuals among medicated patients (*n* = 402). Pearson correlations are displayed, with a fitted linear regression line. 1/f Aperiodic Exponent residuals represent individual 1/f values adjusted for covariates (age, sex, and head motion) using linear regression.

## Discussion

In this multimodal investigation integrating rs-fMRI and EEG across two large, independent cohorts, we found convergent evidence for a shift in cortical excitation-inhibition (E/I) balance toward excitation in psychosis. The rs-fMRI data showed that there were widespread reductions in the HE in people with Early Psychosis. Complementing this, the EEG data pointed to a flattening of the aperiodic 1/f spectral slope in people with chronic psychosis, a pattern similarly associated with reduced inhibitory drive, as previously reported in Schizophrenia (34). Together, these results suggest that E/I imbalance is as a systems-level feature of psychosis, detectable across different neuroimaging modalities and illness stages.

The rs-fMRI findings revealed widespread but not uniform HE reductions. The strongest effects were localized to somatomotor, insular-opercular, midcingulate, dorsolateral prefrontal, and parietal cortices - regions central to sensory integration, cognitive control, and salience processing. Thalamic reductions further highlight a potential disruption in thalamocortical loops, which are consistently implicated in psychosis (35). This is consistent with emerging work on sensorimotor gating deficits (36), somatomotor network dysconnectivity evidence (37), and loss of inhibitory control within thalamo-cortical loops (4). Parallel EEG slope flattening supports multimodal convergence.

These results align with a mechanistic understanding of psychosis rooted in cortical disinhibition. Convergent postmortem (38, 39), animal (40), and computational (19, 41) work implicates deficient GABAergic inhibition, particularly within parvalbumin-pozitive interneurons, as a key driver of microcircuit instability. When inhibitory feedback is compromised, pyramidal neurons exhibit increased baseline firing, high-frequency power becomes accentuated, and the temporal persistence of network activity collapses: precisely the features captured by a reduced HE and flattened 1/f slope (2, 3, 42).

Potassium channels, which critically shape neuronal resting potentials and spike-frequency adaptation, are of particular relevance to psychosis. Dysfunction in these channels can increase baseline excitability while impairing the precision of task-evoked responses. KV3 family channels are most classically linked to parvalabumin-pozitive interneuron physiology (43, 44), and canonical PV-linked KV3 genes were present in the enriched set, although the most prominent association was with KCNB2 (a KV2-family delayed rectifier enriched in pyramidal neurons) (45). This finding suggests that HE may capture regional differences in intrinsic neuronal electrophysiological properties, including those related to pyramidal cell ion-channel expression.

Despite capturing case-control differences, neither HE nor the aperiodic slope robustly predicted the severity of symptoms or cognitive impairments. One interpretation is that, although they index a shared microscale substrate, E/I imbalance represents a stable, trait-like neural vulnerability, while behavioural expression depends on secondary modulatory influences such as dopaminergic tone and network level compensation (46, 47). Alternatively, the mismatch may reflect the distal relationship between microscale neuronal dynamics and clinical phenotypes, confounded by limitations of clinical scales, which show modest reliability (48). Futhermore, medication is likely to shift symptom scores whereas HE abnormalities were robust to antipsychotic medication status and dosage, thereby potentially disrupting any brain-behaviour correlation.

Receptor density mapping further implicated noradrenergic, cholinergic, serotonergic, glutamatergic and dopaminergic systems, of which the noradrenergic and cholinergic markers were the most stable This is consistent with evidence that reduced cholinergic tone weakens inhibitory control. These findings are timely given renewed interest in muscarinic modulation, supported by the clinical efficacy of the M1/M4 agonist xanomeline-trospium (KarXT) in improving psychotic symptoms. Neuroimaging markers of inhibitory tone could help identify individuals most likely to benefit from such interventions (49). Similarly, the identification of potassium-channel enrichment supports ongoing efforts to develop Kv3 agonists as potential treatments for restoring PV interneuron function.

Beyond mechanistic insights, these findings have clear translational potential. Because HE and the aperiodic slope quantify intrinsic, task-free neural dynamics, they are efficient, noninvasive and suitable for longitudinal use. As biomarkers sensitive to inhibitory tone, they could potentially inform early detection and patient stratification. Their applicability across species offers a pathway to mechanistically informed, cross-species biomarker development, an essential step toward precision psychiatry.

This study was limited by its cross-sectional design, which restricted causal inference and prevented modelling of illness-stage effects because time since onset was not available at individual subject level. The HE and aperiodic 1/f spectrum are indirect measures of E/I balance, and further concurrent pharmacological challenge studies would strengthen their biological specificity. Although no relationship was seen with antipsychotic exposure, analyses in medicaton-naive cohorts would be of value. Finally, transcriptomic analyses rely on postmortem samples from a relatively small sample of individuals that may not generalize across populations.

Future studies could validate these markers using multimodal and pharmacological designs. Combining HE and aperiodic slope in the same cohorts with perturbation using E/I modulators could clarify their relationship. Longitudinal work following individuals across illness stages will determine whether these indices track progression or treatment response. Large, harmonized datasets are also needed to test their reliability and clinical utility as biomarkers of cortical E/I imbalance and treatment response prediction.

## Data availability

The data analysed in this study were obtained from the Human Connectome Project–Early Psychosis (HCP-EP) and the Bipolar and Schizophrenia Network on Intermediate Phenotypes 2 (BSNIP-2) datasets. These datasets are available through the NIMH Data Archive (NDA) and/or dbGaP under controlled access and are subject to data use agreements. Derived data supporting the findings of this study are available from the corresponding author upon reasonable request.

## Aknowedgments

Research was funded by the Wellcome Trust (224625/Z/21/Z), Brain and Behaviour Research Foundation (28891), and Academy of Medical Sciences (SGL023\1009). IV, RAM, PM, and DO are supported by the NIHR Oxford Health Biomedical Research Centre. The views expressed are those of the author(s) and not necessarily those of the NIHR or the Department of Health and Social Care.

## Conflict of Interests

IV, ZL have no competing interests to declare. RAM has received speaker/consultancy fees from Angelini Pharma, Boehringer Ingelheim, Bristol Myers Squibb, Janssen, Karuna, Lundbeck, Newron, Otsuka, and Viatris, and co-directs a company that designs digital resources to support treatment of mental ill health. MMN is a Principal Applied Scientist at Microsoft AI.

## References

1. Isaacson Jeffry S, Scanziani M. How Inhibition Shapes Cortical Activity. Neuron. 2011;72(2):231–43.

2. Yizhar O, Fenno LE, Prigge M, Schneider F, Davidson TJ, O’Shea DJ, et al. Neocortical excitation/inhibition balance in information processing and social dysfunction. Nature. 2011;477(7363):171–8.

3. Uhlhaas PJ, Singer W. Abnormal neural oscillations and synchrony in schizophrenia. Nature Reviews Neuroscience. 2010;11(2):100–13.

4. Anticevic A, Lisman J. How Can Global Alteration of Excitation/Inhibition Balance Lead to the Local Dysfunctions That Underlie Schizophrenia? Biological Psychiatry. 2017;81(10):818–20.

5. Menon V. Large-scale brain networks and psychopathology: a unifying triple network model. Trends in Cognitive Sciences. 2011;15(10):483–506.

6. Nour MM, Liu Y, El-Gaby M, McCutcheon RA, Dolan RJ. Cognitive maps and schizophrenia. Trends in Cognitive Sciences. 2025;29(2):184–200.

7. Murray JD, Anticevic A, Gancsos M, Ichinose M, Corlett PR, Krystal JH, et al. Linking Microcircuit Dysfunction to Cognitive Impairment: Effects of Disinhibition Associated with Schizophrenia in a Cortical Working Memory Model. Cerebral Cortex. 2012;24(4):859–72.

8. Liu Y, Ouyang P, Zheng Y, Mi L, Zhao J, Ning Y, et al. A Selective Review of the Excitatory-Inhibitory Imbalance in Schizophrenia: Underlying Biology, Genetics, Microcircuits, and Symptoms. Frontiers in Cell and Developmental Biology. 2021;9.

9. Handest R, Molstrom I-M, Gram Henriksen M, Hjorthøj C, Nordgaard J. A Systematic Review and Meta-Analysis of the Association Between Psychopathology and Social Functioning in Schizophrenia. Schizophrenia Bulletin. 2023;49(6):1470–85.

10. McCutcheon RA, Keefe RSE, McGuire PK. Cognitive impairment in schizophrenia: aetiology, pathophysiology, and treatment. Molecular Psychiatry. 2023;28(5):1902–18.

11. Ahmad J, Ellis C, Leech R, Voytek B, Garces P, Jones E, et al. From mechanisms to markers: novel noninvasive EEG proxy markers of the neural excitation and inhibition system in humans. Translational Psychiatry. 2022;12(1).

12. Cohen Kadosh R. Rethinking excitation/inhibition balance in the human brain. Nature Reviews Neuroscience. 2025;26(8):451–2.

13. Hardstone R, Poil S-S, Schiavone G, Jansen R, Nikulin VV, Mansvelder HD, et al. Detrended Fluctuation Analysis: A Scale-Free View on Neuronal Oscillations. Frontiers in Physiology. 2012;3.

14. Trakoshis S, Martínez-Cañada P, Rocchi F, Canella C, You W, Chakrabarti B, et al. Intrinsic excitation-inhibition imbalance affects medial prefrontal cortex differently in autistic men versus women. eLife. 2020;9.

15. Nishio M, Ellwood-Lowe ME, Woodburn M, McDermott CL, Park AT, Tooley UA, et al. The Development of Neural Inhibition across Species: Insights from the Hurst Exponent. The Journal of Neuroscience. 2025;45(41).

16. Xie K, Royer J, RodriguezLCruces R, Horwood L, Ngo A, Arafat T, et al. Temporal Lobe Epilepsy Perturbs the BrainLWide ExcitationLInhibition Balance: Associations with Microcircuit Organization, Clinical Parameters, and Cognitive Dysfunction. Advanced Science. 2025;12(9).

17. Voytek B, Knight RT. Dynamic Network Communication as a Unifying Neural Basis for Cognition, Development, Aging, and Disease. Biological Psychiatry. 2015;77(12):1089–97.

18. Nishio M, Ellwood-Lowe ME, Woodburn M, McDermott CL, Park AT, Tooley UA, et al. 2024.

19. Gao R, Peterson EJ, Voytek B. Inferring synaptic excitation/inhibition balance from field potentials. NeuroImage. 2017;158:70–8.

20. Glasser MF, Smith SM, Marcus DS, Andersson JLR, Auerbach EJ, Behrens TEJ, et al. The Human Connectome Project’s neuroimaging approach. Nature Neuroscience. 2016;19(9):1175–87.

21. Tamminga CA, Ivleva EI, Keshavan MS, Pearlson GD, Clementz BA, Witte B, et al. Clinical Phenotypes of Psychosis in the Bipolar-Schizophrenia Network on Intermediate Phenotypes (B-SNIP). American Journal of Psychiatry. 2013;170(11):1263–74.

22. Esteban O, Markiewicz CJ, Blair RW, Moodie CA, Isik AI, Erramuzpe A, et al. fMRIPrep: a robust preprocessing pipeline for functional MRI. Nature Methods. 2018;16(1):111–6.

23. Patel AX, Bullmore ET. A wavelet-based estimator of the degrees of freedom in denoised fMRI time series for probabilistic testing of functional connectivity and brain graphs. NeuroImage. 2016;142:14–26.

24. Huang C-C, Rolls ET, Feng J, Lin C-P. An extended Human Connectome Project multimodal parcellation atlas of the human cortex and subcortical areas. Brain Structure and Function. 2021;227(3):763–78.

25. Gramfort A, Luessi M, Larson E, Engemann DA, Strohmeier D, Brodbeck C, et al. MNE software for processing MEG and EEG data. NeuroImage. 2014;86:446–60.

26. Donoghue T, Haller M, Peterson EJ, Varma P, Sebastian P, Gao R, et al. Parameterizing neural power spectra into periodic and aperiodic components. Nature Neuroscience. 2020;23(12):1655–65.

27. Fortin J-P, Cullen N, Sheline YI, Taylor WD, Aselcioglu I, Cook PA, et al. Harmonization of cortical thickness measurements across scanners and sites. NeuroImage. 2018;167:104–20.

28. Thomas Yeo BT, Krienen FM, Sepulcre J, Sabuncu MR, Lashkari D, Hollinshead M, et al. The organization of the human cerebral cortex estimated by intrinsic functional connectivity. Journal of Neurophysiology. 2011;106(3):1125–65.

29. Shen EH, Overly CC, Jones AR. The Allen Human Brain Atlas. Trends in Neurosciences. 2012;35(12):711–4.

30. Hawrylycz MJ, Lein ES, Guillozet-Bongaarts AL, Shen EH, Ng L, Miller JA, et al. An anatomically comprehensive atlas of the adult human brain transcriptome. Nature. 2012;489(7416):391–9.

31. Markello RD, Arnatkeviciute A, Poline J-B, Fulcher BD, Fornito A, Misic B. 2021.

32. Eden E, Navon R, Steinfeld I, Lipson D, Yakhini Z. GOrilla: a tool for discovery and visualization of enriched GO terms in ranked gene lists. BMC Bioinformatics. 2009;10(1).

33. Hansen JY, Shafiei G, Markello RD, Smart K, Cox SML, Nørgaard M, et al. Mapping neurotransmitter systems to the structural and functional organization of the human neocortex. Nature Neuroscience. 2022;25(11):1569–81.

34. Hasanaj G, Jaeger I, Karsli B, Schulz E, Boudriot E, Roell L, et al. Periodic and Aperiodic Alterations of RestingLState EEG in Schizophrenia Spectrum Disorders: Cognitive and Clinical Insights. European Journal of Neuroscience. 2025;62(7).

35. Anticevic A, Halassa MM. The thalamus in psychosis spectrum disorder. Frontiers in Neuroscience. 2023;17.

36. San-Martin R, Castro LA, Menezes PR, Fraga FJ, Simões PW, Salum C. Meta-Analysis of Sensorimotor Gating Deficits in Patients With Schizophrenia Evaluated by Prepulse Inhibition Test. Schizophrenia Bulletin. 2020;46(6):1482–97.

37. Keyvanfard F, Schmid A-K, Nasiraei-Moghaddam A. Functional Connectivity Alterations of Within and Between Networks in Schizophrenia: A Retrospective Study. Basic and Clinical Neuroscience Journal. 2023;14(3):397–410.

38. Lewis DA. Cortical circuit dysfunction and cognitive deficits in schizophrenia – implications for preemptive interventions. European Journal of Neuroscience. 2012;35(12):1871–8.

39. Kaar SJ, Angelescu I, Marques TR, Howes OD. Pre-frontal parvalbumin interneurons in schizophrenia: a meta-analysis of post-mortem studies. Journal of Neural Transmission. 2019;126(12):1637–51.

40. Sohal VS, Zhang F, Yizhar O, Deisseroth K. Parvalbumin neurons and gamma rhythms enhance cortical circuit performance. Nature. 2009;459(7247):698–702.

41. Gonzalez-Burgos G, Cho RY, Lewis DA. Alterations in Cortical Network Oscillations and Parvalbumin Neurons in Schizophrenia. Biological Psychiatry. 2015;77(12):1031–40.

42. Bartos M, Vida I, Jonas P. Synaptic mechanisms of synchronized gamma oscillations in inhibitory interneuron networks. Nature Reviews Neuroscience. 2007;8(1):45–56.

43. Kaar SJ, Nottage JF, Angelescu I, Marques TR, Howes OD. Gamma Oscillations and Potassium Channel Modulation in Schizophrenia: Targeting GABAergic Dysfunction. Clinical EEG and Neuroscience. 2023;55(2):203–13.

44. Faulkner IE, Pajak RZ, Harte MK, Glazier JD, Hager R. Voltage-gated potassium channels as a potential therapeutic target for the treatment of neurological and psychiatric disorders. Frontiers in Cellular Neuroscience. 2024;18.

45. Mohamed ZA, Li J, Wen J, Jia F, Banerjee S. The KCNB2 gene and its role in neurodevelopmental disorders: Implications for genetics and therapeutic advances. Clinica Chimica Acta. 2025;566.

46. Howes O, McCutcheon R, Stone J. Glutamate and dopamine in schizophrenia: An update for the 21stcentury. Journal of Psychopharmacology. 2015;29(2):97–115.

47. McCutcheon RA, Krystal JH, Howes OD. Dopamine and glutamate in schizophrenia: biology, symptoms and treatment. World Psychiatry. 2020;19(1):15–33.

48. Geck S, Roithmeier M, Bühner M, Wehr S, Weigel L, Priller J, et al. COSMIN systematic review and meta-analysis of the measurement properties of the Positive and Negative Syndrome Scale (PANSS). eClinicalMedicine. 2025;82.

49. Kaul I, Sawchak S, Walling DP, Tamminga CA, Breier A, Zhu H, et al. Efficacy and Safety of Xanomeline-Trospium Chloride in Schizophrenia. JAMA Psychiatry. 2024;81(8).

