## Supplemental Material for "Convergent Multimodal Evidence of Cortical Excitation-Inhibition Imbalance in Psychosis"

### I. PART A: fMRI Study Supplemental Material.

#### 1.1 Pre-processing methods detail.

See Fig. 1 for an overview of rs-fMRI methods.

##### 1.1.1 fMRI pre-processing

Neuroimaging data pre-processing was conducted using the fMRIPrep pipeline v25.0.0 using standard settings. Pre-processing of the functional data included steps such as slice time correction via 3dTshift from AFNI (1), motion correction via MCFLIRT (2), co-registration of BOLD images to the T1-weighted anatomical using boundary-based registration (3) via FreeSurfer bbregister command and spatial normalization to MNI152 space, which are all concatenated in one step via antsApplyTransforms. Framewise displacement was computed for each run to quantify motion (4).

For the structural MRI, the T1-weighted (T1w) images were corrected for intensity non-uniformity using **N4BiasFieldCorrection** (5), distributed with Advanced Normalization Tools (ANTs)2.5.4 (6) and skull-stripped with the  **antsBrainExtraction.sh** using the **OASIS** template. **FreeSurfer** 7.3.2 (7, 8) was employed for surface reconstruction via the recon-all command. The brain mask obtained during skull-stripping was further refined using a method inspired by **Mindboggle** to reconcile segmentations derived from both **ANTs** and **FreeSurfer (9).** Spatial normalization to the **MNI152NLin2009cAYM (10)** was performed via nonlinear registration using the antsRegistration tool. Finally, brain tissue segmentation into grey matter (GM), white matter (WM), and cerebrospinal fluid (CSF) was performed using **FAST** (FSL) (11).

##### 1.1.2 fMRI additional denoising

Additional denoising consists of wavelet despiking (BrainWavelet toolbox v2.0, threshold = 10) (12) and 13 parameter nuisance regression (6 motion parameters, their first derivatives and cerebrospinal fluid signal) (13). The wavelet despiking method decomposes each voxel’s BOLD signal into multiple frequency scales using discrete wavelet transforms, identifies non-Gaussian transients (spikes) at each scale, and suppresses them adaptively while preserving the underlying neural signal. This approach reduces high-frequency noise without excessive temporal smoothing, improving signal-to-noise ratio and the reliability of derived estimates (12).

##### 1.1.3 fMRI parcellation

Parcel-wise fMRI time series were extracted using the HCPex atlas (14) a volumetric extension of the Human Connectome Project Multi-Modal Parcellation (15). HCPex comprises 360 cortical parcels derived from the original surface-based atlas, adapted for use with volumetric neuroimaging software, and includes an additional 66 subcortical regions (33 per hemisphere). For each parcel, BOLD signal was averaged across all voxels within AFNI-derived brain masks; voxels outside the mask were excluded. Per-parcel voxel coverage was recorded for quality control.

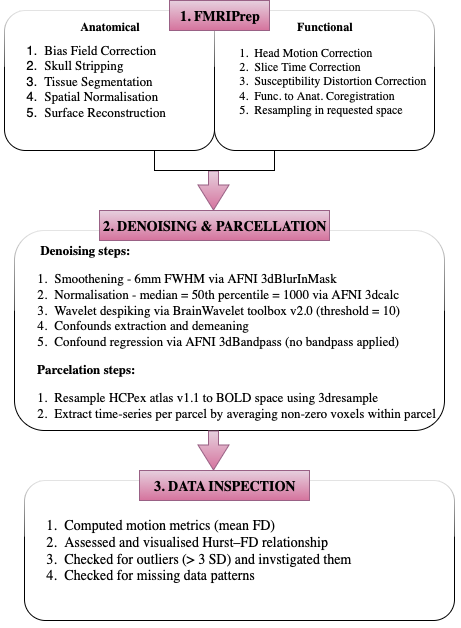

**Figure 1.** Visual representation of the Pre-processing and Denoising Steps.

##### 1.1.4 fMRI site harmonisation

Regional Hurst exponent (HE) and cortical thickness (CT) measures were processed independently but using an identical harmonisation pipeline employing ComBat (16). For each modality, parcel-wise values were first assembled across subjects and then harmonised separately using the same procedure, with site specified as the batch variable and age, sex, and diagnosis included as covariates of interest. Harmonisation was performed independently for HE and CT to account for modality-specific noise characteristics, after which whole-brain summary measures were derived and downstream analyses conducted on the harmonised data.

#### 1.2 Quality checks before analysis

All pre-processing outputs were visually inspected to confirm accurate brain extraction, tissue segmentation, and spatial alignment of functional and anatomical images. Subjects with mFD exceeding 0.25 mm were excluded. To investigate whether motion may influence the hurst exponent (HE), we conducted linear regression analyses within groups, modelling whole brain HE as a function of mean framewise displacement (mFD).The within group analysis was required to avoid Simpson’s paradox (17). The whole brain HE and mFD association was negligible (*β* = –0.08, *P* = 0.21) (Fig. 2).

Similarly, to identify potential outliers without introducing group-level bias, we computed z-scores for the whole-brain HE separately within each diagnosis group. Subjects with *|z|* > 3 SD were flagged as potential outliers, striking a balance between sensitivity and specificity (18). Outliers > 3SD but not cofounded by motion were kept for sensitivity analysis (Fig. 2). In our analysis we only identified one healthy control outlier, which we excluded for our final sample size and included in sensitivity analysis. See results in supplementary section 1.5.

Finally, regional HEs values were inspected for missing data both visually, using binary heatmaps, and quantitatively, by calculating the proportion of missing values per region and per subject. In our final data set, after excluding the outlier, we only had missing values for one parcel value for 1 subject, requiring no imputation before further analysis as per established guidelines (19).

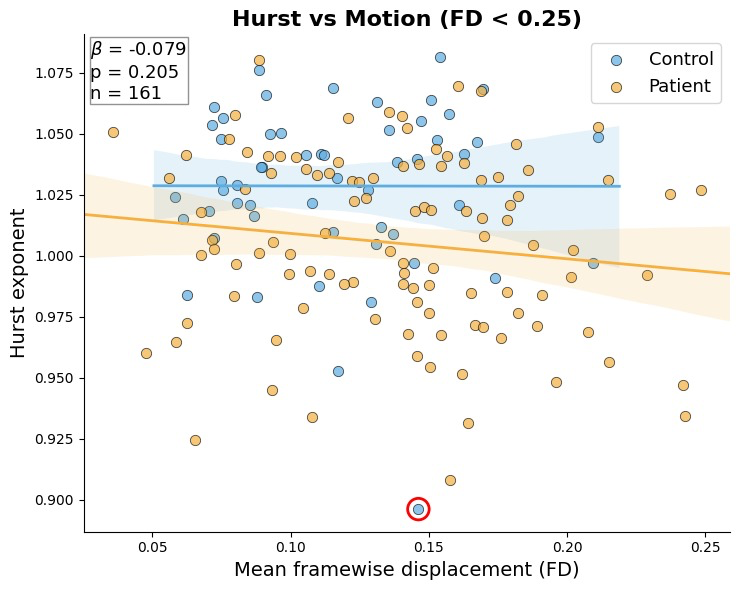

**Figure 2.** Scatter plot showing the relationship between the **whole brain HE** and **mFD,** for participants with mFD < 0.25. Each dot represents an individual subject, with **blue dots** indicating control participants and **orange dots** indicating patient participants. Solid lines show linear regression fits for each group, with shaded regions representing the 95% confidence intervals. The inset box reports the regression slope (*β*), p-value, and total sample size (*n*). The **red-circled dot** highlights a control outlier captured by the |z| > 3SD rule.

#### 1.3 Individual analysis methods overview

##### 1.3.1 Network enrichment

For each cortical parcel, ordinary least squares (OLS) regression was used to estimate the effect of diagnostic group (Control vs Patient) on the variable of interest, controlling for age, sex, and mFD. The resulting parcel-wise β coefficients were then summarized at the level of large-scale Yeo-7 functional networks using surface-based fractional overlaps between the HCPex parcellation and the Yeo-7 atlas (20). Network-level effects (Δ) were computed as the difference between each network’s surface-weighted mean β and the corresponding global (whole-cortex) surface-weighted mean β:

$$\Delta_{n}=\bar{\beta}_{n}^{\left( weighted \right)}-\bar{\beta}_{global}^{\left( weighted \right)}.$$

Weights corresponded to the proportion of each parcel’s cortical surface area overlapping each Yeo-7 network. To assess statistical significance, a non-parametric permutation test (10,000 iterations) was performed by randomly permuting the phenotype labels and recomputing the parcel-wise OLS fits and resulting Δ values for each permutation. Two-sided empirical *p*-values were derived as the proportion of permuted |Δ| values exceeding the observed |Δ|.

##### 1.3.2 Cortical thickness

For each cortical parcel, an ordinary least squares (OLS) regression model was fitted separately for HE and CT data using the formula Y ∼ Phenotype + Age + Sex + MeanFD. The regression coefficient (β) for the Phenotype term was extracted from each parcel, yielding vectors of β values across the cortex for both modalities. Cross-modal association between HE and CT was quantified using Pearson (r) correlations of these β vectors. Additionally, the associations between whole brain CT and whole brain HE across subjects were assessed using Pearson correlation. To assess statistical significance, a non-parametric permutation test (10,000 iterations) was performed by randomly permuting the phenotype labels, with full model refitting at each iteration.

##### 1.3.3 Gene expression analysis pipeline

**Abagen Pipeline**. Regional microarray expression data were obtained from 6 post-mortem brains (1 female, ages 24.0–57.0, 42.50 +/- 13.38) provided by the Allen Human Brain Atlas (AHBA, <https://human.brain-map.org>) (21). Since the AHBA included only two subjects with right sided samples, limiting the data on the right side, keeping with similar analyses, we only consider left hemisphere samples. Data were processed with the abagen toolbox (version 0.1. 4+15.gdc4a007.dirty; <https://github.com/rmarkello/abagen>) (22) using the HCPEX v1.1 volumetric atlas in MNI space (<https://github.com/wayalan/HCPex>) (14, 23).

First, microarray probes were reannotated using data provided by Arnatkevic̆iūtė et al. (24); probes not matched to a valid Entrez ID were discarded. Next, probes were filtered based on their expression intensity relative to background noise (25), such that probes with intensity less than the background in >=50.00% of samples across donors were discarded, yielding 31,569 probes. When multiple probes indexed the expression of the same gene, we selected and used the probe with the most consistent pattern of regional variation across donors (i.e., differential stability) (26), calculated with:

$$\Delta_{S}(p)=\frac{1}{\binom{N}{2}} \sum_{i=1}^{N-1} \sum_{j=i+1}^{N} \rho[B_{i}(p),B_{j}(p)]$$

where $\rho$ is Spearman’s rank correlation of the expression of a single probe, p, across regions in two donors $B_{i}$ and $B_{j}$, and N is the total number of donors. Here, regions correspond to the structural designations provided in the ontology from the AHBA.

The MNI coordinates of tissue samples were updated to those generated via non-linear registration using the Advanced Normalization Tools (ANTs; https://github.com/chrisfilo/alleninf). Samples were assigned to brain regions in the provided atlas if their MNI coordinates were within 2 mm of a given parcel. If a brain region was not assigned a sample from any donor based on the above procedure, the tissue sample closest to the centroid of that parcel was identified independently for each donor. The average of these samples was taken across donors, weighted by the distance between the parcel centroid and the sample, to obtain an estimate of the parcellated expression values for the missing region. This procedure was performed for 64 regions that were not assigned tissue samples. All tissue samples not assigned to a brain region in the provided atlas were discarded.

Inter-subject variation was addressed by normalizing tissue sample expression values across genes using a robust sigmoid function (27):

$$x_{norm}=\frac{1}{1+exp(-\frac{(x-\langle x\rangle)}{\text{IQR}_{x}})}$$

where $\langle x\rangle$ is the median and $\text{IQR}_{x}$ is the normalized interquartile range of the expression of a single tissue sample across genes. Normalized expression values were then rescaled to the unit interval:

$$x_{scaled}=\frac{x_{norm}-min(x_{norm})}{max(x_{norm})-min(x_{norm})}$$

Gene expression values were then normalized across tissue samples using an identical procedure. Samples assigned to the same brain region were averaged separately for each donor and then across donors, yielding a regional expression matrix with rows corresponding to cortical brain regions from the left hemisphere (*n* = 180) and columns corresponding to the retained genes for downstream analysis (*n* = 15.633).

**PLSR Pipeline**. Partial Least Square Regression (PLSR) was used as a multivariate association method to relate high-dimensional parcel-wise gene expression data to phenotype-associated variation in HE. Cross-validation (CV) was used to control model complexity and assess out-of-sample association strength. All PLSR analyses were performed using z-scored predictors and targets, with scaling parameters estimated only within training folds and applied to held-out folds.

The PLSR pipeline comprised two steps: (i) a primary nested cross-validation model for component selection and test of statistical significance, and (ii) a final cross-validated parsimonious model with a fixed number of latent variables (LV) informed by the initial model, for interpretation.

Statistical significance was assessed using nonparametric permutation testing (*n* = 10,000 iterations). For each permutation, case-control labels were randomly reassigned across subjects, and the OLS model was re-estimated to generate a permuted parcel-wise phenotype-effect map. The full PLSR analysis and CV, was then repeated using this permuted Y map, yielding a null distribution of cross-validated association metrics (R² and Pearson’s r).

*Primary model:*

The primary PLSR analysis employed nested cross-validation. An outer 10-fold CV loop was used to generate out-of-fold (OOF) estimates, while an inner 5-fold CV loop was used to select the number of latent variables (k) based on R² performance. Inner CV results were summarized within each outer fold, and the mean and standard error of these results were then computed across all outer folds to apply a global 1-SE rule (28), selecting the smallest k within one standard error of the maximum. Cross-validated R² and Pearson’s r, computed from OOF estimates across outer folds, were used as measures of out-of-sample association strength. The selected number of LVs informed the final model specification. The results of this model are reported in the main manuscript and the k selection by 1-SE rule plotting can be visualised in the supplement below at section 1.4 Supplementary analyses results.

*Final Parsimonious model:*

Using the latent variable dimensionality selected in the primary analysis, the model was refit with an identical specification and identical CV splits, the only difference being no further nested hyperparameter optimisation. This fixed-complexity refit was used to characterise and interpret the dominant, stable covariance structure underlying the primary association. Variable importance in projection (VIP) scores were computed from this parsimonious fixed-dimensional model to characterise the contribution of individual predictors to the stable covariance structure identified in the primary analysis.

**Gene Enrichment.** Gene-wise contribution was quantified with Variable Importance in Projection (VIP). Uncertainty in each gene’s VIP was estimated by bootstrap resampling of parcels (left-hemisphere cortical parcels; *B* = 10,000), refitting the PLSR on each bootstrap sample. We standardized each gene’s VIP by its bootstrap-derived standard error to obtain a z score and ranked genes by this standardized influence score. To characterize biology while avoiding inflation by broadly brain-expressed genes, we first restricted the VIP-ranked list to transcripts with nonzero RNA expression (Transcripts Per Million (TPM) >0) in cerebral cortex according to the Human Protein Atlas/GTEx brain dataset (29) (https://www.proteinatlas.org/download/tsv/rna_brain_gtex.tsv.zip). The cortex-refined, rank-ordered list was submitted to GOrilla (30) (ranked-list mode, separately in ascending and descending order). Gene Ontology terms were deemed enriched at FDR-corrected *q* < 0.05 with a minimum of three overlapping genes.

##### 1.3.4 Receptor density maps analysis

We analysed neurotransmitter receptor density maps compiled by Hansen et al. (31) (19 targets). Volumetric PET/SPECT maps in MNI152 space were parcellated to the HCPex cortical atlas with neuromaps’ Parcellater; where multiple tracers existed for a target, maps were z-scored and combined by sample-size weighted averaging to yield one parcel-wise map per receptor. The parcel-wise outcome was the OLS phenotype coefficient (*β* for Phenotype in: HE ~ Phenotype + Age + Sex + FD), aligned to the same parcels. We then applied the same PLSR pipeline described above. To summarize receptor contributions, we computed VIP and quantified stability via bootstrap resampling of parcels (*B* = 10,000); these VIP estimates were used to interpret relative receptor importance in the PLSR model.

##### 1.3.5 Clinical and cognitive PLSR models

Clinical and cognitive outcomes were related to regional HE features using the same PLSR pipeline described above. Parcel-wise cortical HE values served as the input feature matrix, and outcomes were either multivariate (individual PANSS items) or scalar (PANSS total scores or a cognitive composite). See Table 1.

| **Model** | **X (n features)** | **y (n outcomes)** |
| --- | --- | --- |
| **Clinical (PANSS)** | | |
| 1 | PANSS Subscales (3) | HA (1) |
| 2 | PANSS All items (30) | HA (1) |
| 3 | PANSS All items (30) | HE parcels (360) |
| 4 | HE parcels (360) | PANSS All items (30) |
| 5 | PANSS Subscales (3) | HE parcels (360) |
| 6 | HE parcels (360) | PANSS Subscales (3) |
| 7 | HE parcels (360) | PANSS Negative (1) |
| **Cognition** | | |
| 8 | HE parcels (360) | Cognitive composite score (1) |

**Table 1**. Clinical and Cognitive PLSR Models.

#### 1.4 Supplementary analyses results

##### 1.4.1 HE case - control differences top 20 contributing cortical and subcortical parcels.

| **Parcel** | ***β* Phenotype** | ***t* value** | ***P* value** | ***q* FDR** |
| --- | --- | --- | --- | --- |
| SomatosensoryandMotor_3a_Right | 0.091 | 5.829 | <0.001 | <0.001 |
| ParacentralLobularandMidCingulate_24dd_Right | 0.073 | 4.041 | <0.001 | 0.002 |
| InsularandFrontalOpercular_PoI2_Right | 0.072 | 4.268 | <0.001 | 0.002 |
| ParacentralLobularandMidCingulate_5m_Right | 0.071 | 4.331 | <0.001 | 0.002 |
| ParacentralLobularandMidCingulate_24dd_Left | 0.071 | 4.187 | <0.001 | 0.002 |
| PosteriorOpercular_OP2-3_Left | 0.070 | 3.565 | <0.001 | 0.005 |
| EarlyAuditory_RI_Right | 0.068 | 3.708 | <0.001 | 0.004 |
| InsularandFrontalOpercular_Ig_Left | 0.068 | 3.763 | <0.001 | 0.004 |
| InsularandFrontalOpercular_FOP2_Left | 0.068 | 3.963 | <0.001 | 0.003 |
| InsularandFrontalOpercular_Ig_Right | 0.068 | 3.776 | <0.001 | 0.004 |
| ParacentralLobularandMidCingulate_5m_Left | 0.068 | 3.929 | <0.001 | 0.003 |
| PosteriorOpercular_OP4_Left | 0.068 | 4.233 | <0.001 | 0.002 |
| SuperiorParietal_LIPd_Right | 0.068 | 3.722 | <0.001 | 0.004 |
| SomatosensoryandMotor_3a_Left | 0.066 | 4.181 | <0.001 | 0.002 |
| PosteriorOpercular_OP2-3_Right | 0.065 | 3.235 | 0.001 | 0.010 |
| SomatosensoryandMotor_3b_Right | 0.065 | 4.504 | <0.001 | 0.002 |
| Premotor_PEF_Right | 0.064 | 4.190 | <0.001 | 0.002 |
| ParacentralLobularandMidCingulate_6mp_Left | 0.062 | 3.502 | 0.001 | 0.005 |
| InferiorFrontal_IFJp_Left | 0.062 | 3.819 | <0.001 | 0.004 |
| PosteriorOpercular_43_Left | 0.061 | 4.058 | <0.001 | 0.002 |

**Table 2A.** Top 20 cortical parcels showing the largest case–control differences in Hurst exponent (HE). Results from parcel-wise ordinary least squares (OLS) models of HE ~ Phenotype + Age + Sex + MeanFD, where the Phenotype term represents the case–control contrast (Control – Patient). Columns show the standardized regression coefficient (*β*), corresponding t-value, nominal p-value, and FDR-corrected q-value. Positive β indicates lower HE in patients relative to controls. The strongest effects were observed in somatosensory, mid-cingulate, and insular/frontal-opercular regions.

| **Parcel** | ***β* Phenotype** | ***t* value** | ***P* value** | ***q* FDR** |
| --- | --- | --- | --- | --- |
| Thalamic_Nuclei_VLa_Right | 0.051 | 3.871 | 0.000 | 0.004 |
| Thalamic_Nuclei_VLp_Left | 0.044 | 3.152 | 0.002 | 0.012 |
| Thalamic_Nuclei_VLp_Right | 0.043 | 3.129 | 0.002 | 0.012 |
| Thalamic_Nuclei_LD_Left | 0.040 | 4.246 | 0.000 | 0.002 |
| Thalamic_Nuclei_LGN_Left | 0.038 | 3.500 | 0.001 | 0.005 |
| Thalamic_Nuclei_VA_Right | 0.038 | 3.144 | 0.002 | 0.012 |
| Thalamic_Nuclei_VPL_Left | 0.035 | 2.837 | 0.005 | 0.021 |
| Thalamic_Nuclei_PuM_Left | 0.035 | 2.518 | 0.013 | 0.034 |
| Thalamic_Nuclei_AV_Right | 0.034 | 2.952 | 0.004 | 0.017 |
| Thalamic_Nuclei_VPL_Right | 0.034 | 2.809 | 0.006 | 0.022 |
| Thalamic_Nuclei_LP_Right | 0.033 | 3.131 | 0.002 | 0.012 |
| Thalamic_Nuclei_PuM_Right | 0.033 | 2.293 | 0.023 | 0.050 |
| Thalamic_Nuclei_VLa_Left | 0.032 | 2.412 | 0.017 | 0.043 |
| Thalamic_Nuclei_MDl_Left | 0.031 | 2.488 | 0.014 | 0.036 |
| Thalamic_Nuclei_PuI_Left | 0.031 | 2.806 | 0.006 | 0.022 |
| Thalamic_Nuclei_CL_Left | 0.030 | 3.185 | 0.002 | 0.011 |
| Thalamic_Nuclei_MDm_Right | 0.030 | 2.448 | 0.015 | 0.040 |
| Thalamic_Nuclei_CM_Right | 0.029 | 2.773 | 0.006 | 0.022 |
| Thalamic_Nuclei_PuA_Left | 0.028 | 2.457 | 0.015 | 0.039 |
| Thalamic_Nuclei_LP_Left | 0.027 | 2.364 | 0.019 | 0.046 |

**Table 2B.** Top 20 subcortical parcels showing the largest case–control differences in Hurst exponent (HE). Results from parcel-wise ordinary least squares (OLS) models of HE ~ Phenotype + Age + Sex + MeanFD, where the Phenotype term represents the case–control contrast (Control – Patient). Columns show the standardized regression coefficient (*β*), corresponding t-value, nominal p-value, and FDR-corrected q-value. Positive β indicates lower HE in patients relative to controls. The strongest effects were observed bilaterally in thalamus regions.

##### 1.4.2. Regional cortical thickness results

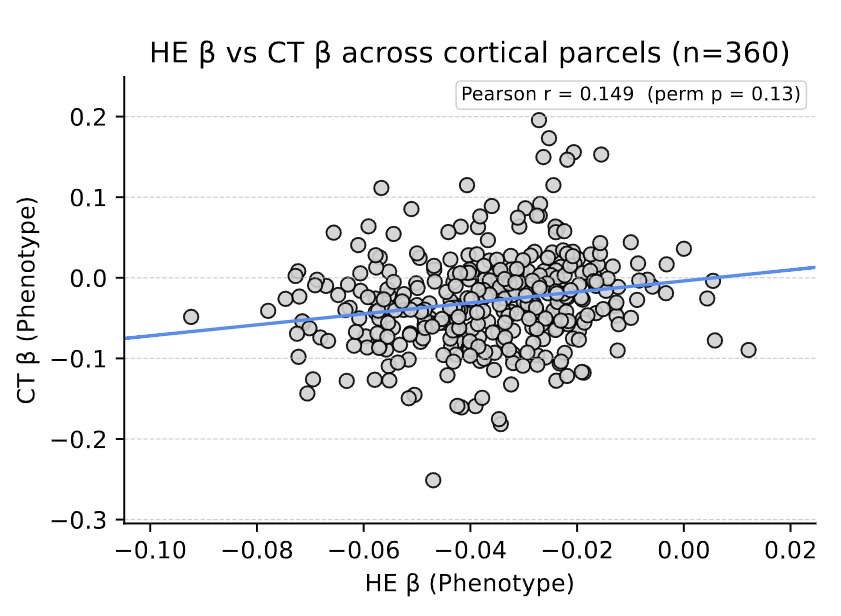

**Figure 3.** Association between HE and CT effects across cortical parcels (*n* = 360). Each point is a parcel. Axes show the OLS β for the Phenotype term from parcel-wise models (HE or CT ~ Phenotype + Age + Sex + MeanFD). The line is the least-squares fit across parcels. We observe a weak positive association: Pearson r = 0.149 (perm *P* = 0.13); where permutation p-values are obtained by shuffling phenotype labels across subjects and recomputing parcel-wise βs before correlating. Positive values indicate parcels where a stronger phenotype effect on HE tends to coincide with a stronger Phenotype effect on CT.

##### 1.4.3 Receptor density PLSR results.

1.4.3.1 Latent variable (*k*) selection in primary model for receptor density

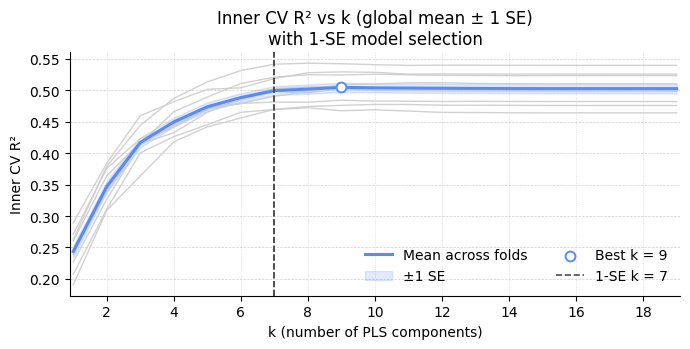

**Figure 4.** Inner cross-validation tuning of PLS components (global mean ±1 SE). Across outer folds, inner CV R² is plotted as a function of the number of PLS components (*k*). Thin grey lines show per-fold tuning curves; the blue line is the mean across folds with a shaded ±1 SE band. The open circle marks the global best by mean CV R² (*k* = 9). Using the 1-SE rule, we selected the smallest k within 1 SE of the best, yielding *k* = 7 (vertical dashed line) for the parsimonious model.

##### 1.4.4 Gene expression results.

1.4.4.1 Gene Expression Primary Model PLSR Results

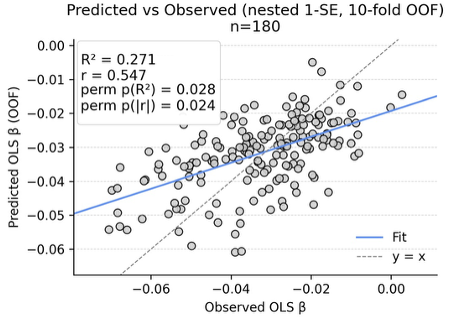

**Figure 5.** Cross-validated association between AHBA gene-expression PLSR estimates and parcel-wise HE phenotype effects (10-fold CV). Out-of-fold model-derived estimates from a PLSR relating Allen Human Brain Atlas gene expression to parcel-wise OLS β (predicted) are plotted against the observed β across left-hemisphere cortical parcels (grey dots, *n* = 180). The solid blue line shows the fitted linear relationship between observed and model-derived values, and the dashed line denotes the identity line (*y* = *x*). Cross-validated association strength is quantified as *R²* = 0.27 (perm *P* = .02) and Pearson’s *r* = 0.54 (perm *P* = .02). Empirical significance was assessed using phenotype-label permutation testing, with parcel-wise *β* estimation and the full CV pipeline recomputed at each permutation.

1.4.4.2 Latent variable (*k*) selection in primary model PLSR for gene expression

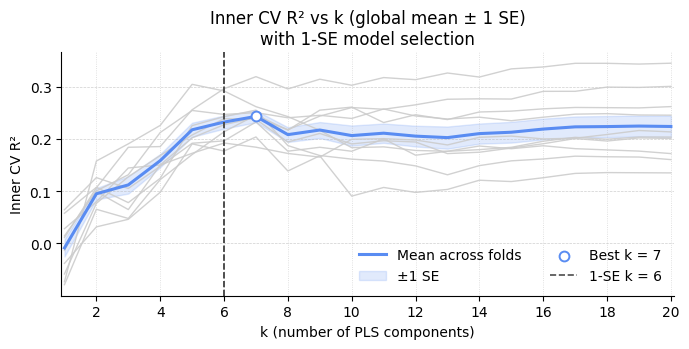

**Figure 6.** Inner cross-validation tuning for gene-expression PLSR (global mean ±1 SE). Inner CV R² is shown as a function of PLSR components (*k*) across outer folds. Thin grey traces are per-fold curves; the blue curve is the mean across folds with a shaded ±1 SE band. The open circle marks the global best by mean CV R² (*k* = 7). Applying the 1-SE rule (choose the smallest k within 1 SE of the best) yields *k* = 6 (vertical dashed line), which was used for the parsimonious model.

1.4.4.3	Correlation between KCNB2 expression and illness effects (Beta Phenotype from HE OLS model)

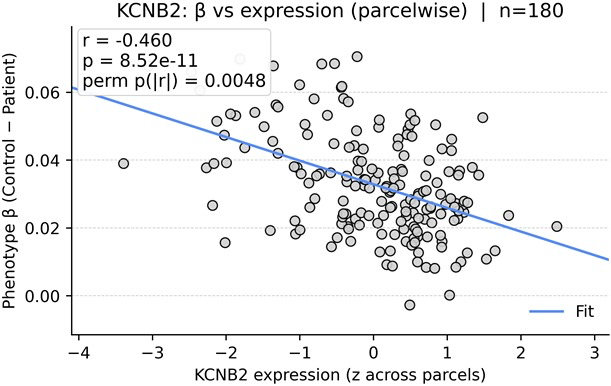

**Figure 7**. KCNB2 expression vs phenotype (parcel-wise). Each dot is a left-hemisphere HCPex parcel. X = KCNB2 expression (z-scored across parcels) and Y = OLS phenotype coefficient (HE ~ Phenotype + Age + Sex + FD). The blue line is the least-squares fit, illustrating the parcel-wise association.

#### 1.5 HE Sensitivity analyses results

| Analysis | Main result *n* = 160  (53 HC, 107 FEP) | Sensitivity *n* = 161  (54 HC, 107 FEP) | Change / Robustness |
| --- | --- | --- | --- |
| Whole Brain Group Differences HC - FEP | *β* = 0.031 (perm *P* < 0.001)  95% CI = [0.015, 0.047] | *β* = 0.028 (perm *P* < 0.001)  95% CI = [0.011, 0.044] | Robust, similar result, remains significant. |
| Regional Hurst group differences | 223/360 cortical regions with *P* < 0.05, 117 with FDR *q* < 0.05  29/66 subcortical regions with *P* < 0.05, 22 with FDR *q* < 0.05 | 181/360 cortical regions with *P* < 0.05, 100 with FDR *q* < 0.05  23/66 subcortical regions with *P* < 0.05, 12 with FDR *q* < 0.05 | Robust, similar result, with most important regions unchanged. |
| Functional Network Enrichment (Yeo) | Somatomotor *Δβ* = 0.019 (perm *P* = 0.001, FDR *q* < 0.05). Default *Δβ* = − 0.007 (perm *P* < 0.05, FDR *q* > 0.05) | Somatomotor *Δβ* = 0.018 (perm *P* < 0.05, FDR *q* < 0.05). Default *Δβ* = − 0.007 (perm P < 0.05, FDR *q* > 0.05) | Robust, similar result, remains significant. |
| Receptor Density Maps PLSR | 1-SE K selection model:  *R²* =0.474 (perm *P* = 0.013)  *r* =0.690 (perm *P* = 0.013)  Parsimonious model (*k*=7):  *R²* = 0.516  *r* = 0.719 (*P* = 1.34e-59)  VIP > 1: NET, VAChT, 5HT4, NMDA, 5HT2a, DAT | 1-SE K selection model:  *R²* = 0.413 (perm *P* = 0.035)  *r* = 0.647 (perm *P* = 0.033)  Parsimonious model (*k*=7):  *R² =* 0.476  *r* = 0.692 (*P* = 1.34e-52)  VIP > 1: NET, VAChT, 5HT4, NMDA, 5HT2a, DAT | Robust, similar result, remains significant |
| Regional Gene Expression PLSR | 1-SE K selection model:  *R² =* 0.271 (perm *P* = 0.028)  *r* = 0.547 (perm *P* = 0.024)  Parsimonious model (*k*=6):  *R²* = 0.321  *r* = 0.582 (*P* = 1.04e-17) | 1-SE K selection model:  *R²* = 0.251 (perm *P* = 0.035)  *r* = 0.523 (perm *P* = 0.035)  Parsimonious model (*k*=6):  *R²* = 0.268  *r* = 0.543 (*P* = 3.5e-15) | Robust, similar result, remains significant |

**Table 3A.**

| Analysis | Main result *n* = 145  (45 HC, 100 FEP) | Sensitivity *n* = 147  (47 HC, 100 FEP) | Change / Robustness |
| --- | --- | --- | --- |
| Regional Cortical Thickness vs Hurst Exponent correlation | Pearson *r* = 0.149 (*P* = 0.005, perm *P* = 0.132) | Pearson *r* = 0.149 (*P* = 0.005, perm *P* = 0.131) | Robust, similar result, remains insignificant. |

**Table 3B.**

**Table 3.** Sensitivity analyses including the previously excluded outlier participant. Sensitivity analyses were conducted to assess the robustness of all primary findings after re-including one participant excluded for being a univariate outlier (>3 SD from the mean Hurst value). Results were compared with the main analyses across all models, including group-level, regional, and multivariate associations. The sensitivity analyses produced highly similar results to the main findings, with all key effects remaining in the same direction and of comparable magnitude. No substantive changes in significance or interpretation were observed.

### II. PART B EEG: Supplementary Material

#### 2.1 Pre-processing methods

EEG pre-processing was performed in Python 3.6 using MNE-Python (32). NeuroScan.cnt EEG recordings marked with ‘eyes open’ and ‘rest’ were imported, and template electrode positions were assigned using a 10–10 montage. Channels were then re-referenced to the common average reference to reduce reference bias after removal of linked mastoid reference electrodes. 0.5–40 Hz band-pass filtering was applied using a zero-phase Hamming-window finite impulse response design (33). Because the low-pass cutoff was 40 Hz, mains-frequency line noise lay outside the analysed band, so notch filtering was not applied. Bad channels were detected using PREP-like metrics (flat channels via ~0.5 µV RMS floor, high-RMS outliers, and low correlation to a robust reference) and, when electrode positions were available, interpolated via spline interpolation with nearest neighbour fallback if spline failed (34). If >30% of EEG channels are bad before interpolation, the run is rejected outright.

Artefacts were removed using an independent component analysis (ICA). Data were high pass filtered at 1 Hz for decomposition, and ICA was attempted using Picard and fallback to Infomax and FastICA. Components strongly correlated with ocular, cardiac, or showing high-frequency/muscle or kurtotic features were excluded, with a maximum of 40% of components removed (35, 36). Cleaned data were resampled to 250 Hz, segmented into 2-second epochs, and epochs exceeding 150 µV peak-to-peak were rejected. Power spectral density (PSD) was computed for each epoch and channel using Welch’s method (1-second windows, 50 % overlap) and interpolated onto a 0.5-Hz grid spanning 1–40 Hz (37).

#### 2.2 1/f analysis

Within each EEG recording, epochs were averaged to yield a run-level mean spectrum, and multiple runs for the same participant were then averaged to produce a subject-level mean PSD. Aperiodic 1/f components of the EEG spectra were then modelled using the FOOOF/SpecParam algorithm (38). For each subject-level PSD, the power spectrum was fitted between 1–40 Hz (within a broader frequency window of 0.5–45 Hz) using a three-parameter knee aperiodic model. This model separates the aperiodic 1/f slope exponent and offset from any periodic oscillatory peaks, providing estimates of underlying scale-free activity. The model parameters (offset, knee, exponent) and goodness-of-fit statistics (R² and error) were extracted for each participant. A combined summary table was generated containing all parameter estimates across the cohort.

Quality control was performed on the spectral fits to ensure physiological plausibility and model reliability. Fits were retained only if the coefficient R² exceeded 0.8, exponents fell between 0 and 5, and offsets ranged from -30 to +30 dB. Any spectra failing these criteria were removed. These quality-filtered outputs formed the basis for all subsequent statistical analyses of 1/f parameters. Automated QC excluded poor-fit or outlier spectra (R² ≥ 0.8, 0 ≤ exponent ≤ 5, −30 ≤ offset ≤ 30).

#### 2.3 1/F spectrum additional results

##### 2.3.1 1/f aperiodic offset case – control differences.

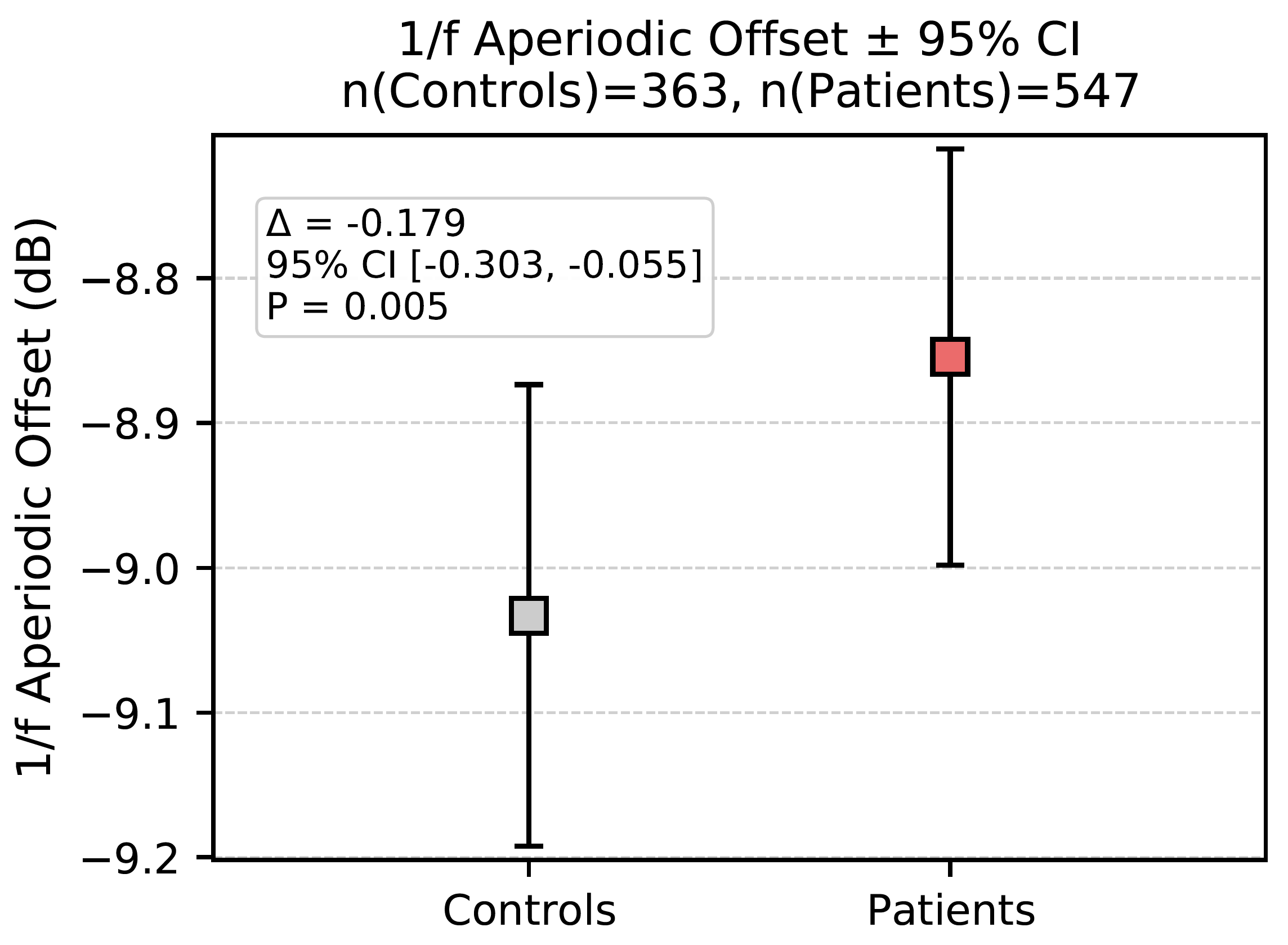

**Figure 8.** 1/f aperiodic offset in healthy controls and psychosis patients. OLS model (offset ~ group + age + sex + exponent + site) demonstrates significantly elevated offsets in psychosis vs healthy controls (*β* = -0.179 ± 0.063, *z* = -2.823, *P* = 0.0049).

**REFERENCES**

1.	Cox RW. AFNI: Software for Analysis and Visualization of Functional Magnetic Resonance Neuroimages. Computers and Biomedical Research. 1996;29(3):162-73.

2.	Jenkinson M. Improved Optimization for the Robust and Accurate Linear Registration and Motion Correction of Brain Images. NeuroImage. 2002;17(2):825-41.

3.	Greve DN, Fischl B. Accurate and robust brain image alignment using boundary-based registration. NeuroImage. 2009;48(1):63-72.

4.	Power JD, Mitra A, Laumann TO, Snyder AZ, Schlaggar BL, Petersen SE. Methods to detect, characterize, and remove motion artifact in resting state fMRI. NeuroImage. 2014;84:320-41.

5.	Tustison NJ, Avants BB, Cook PA, Yuanjie Z, Egan A, Yushkevich PA, et al. N4ITK: Improved N3 Bias Correction. IEEE Transactions on Medical Imaging. 2010;29(6):1310-20.

6.	Avants B, Tustison NJ, Song G. Advanced Normalization Tools: V1.0. The Insight Journal. 2009.

7.	Dale AM, Fischl B, Sereno MI. Cortical Surface-Based Analysis. NeuroImage. 1999;9(2):179-94.

8.	Reuter M, Schmansky NJ, Rosas HD, Fischl B. Within-subject template estimation for unbiased longitudinal image analysis. NeuroImage. 2012;61(4):1402-18.

9.	Schneidman D, Klein A, Ghosh SS, Bao FS, Giard J, Häme Y, et al. Mindboggling morphometry of human brains. PLOS Computational Biology. 2017;13(2).

10.	Fonov VS, Evans AC, McKinstry RC, Almli CR, Collins DL. Unbiased nonlinear average age-appropriate brain templates from birth to adulthood. NeuroImage. 2009;47.

11.	Zhang Y, Brady M, Smith S. Segmentation of brain MR images through a hidden Markov random field model and the expectation-maximization algorithm. IEEE Transactions on Medical Imaging. 2001;20(1):45-57.

12.	Patel AX, Bullmore ET. A wavelet-based estimator of the degrees of freedom in denoised fMRI time series for probabilistic testing of functional connectivity and brain graphs. NeuroImage. 2016;142:14-26.

13.	Trakoshis S, Martínez-Cañada P, Rocchi F, Canella C, You W, Chakrabarti B, et al. Intrinsic excitation-inhibition imbalance affects medial prefrontal cortex differently in autistic men versus women. eLife. 2020;9.

14.	Huang C-C, Rolls ET, Feng J, Lin C-P. An extended Human Connectome Project multimodal parcellation atlas of the human cortex and subcortical areas. Brain Structure and Function. 2021;227(3):763-78.

15.	Glasser MF, Coalson TS, Robinson EC, Hacker CD, Harwell J, Yacoub E, et al. A multi-modal parcellation of human cerebral cortex. Nature. 2016;536(7615):171-8.

16.	Fortin J-P, Cullen N, Sheline YI, Taylor WD, Aselcioglu I, Cook PA, et al. Harmonization of cortical thickness measurements across scanners and sites. NeuroImage. 2018;167:104-20.

17.	Norton HJ, Divine G. Simpson's Paradox … and How to Avoid it. Significance. 2015;12(4):40-3.

18.	Berger A, Kiefer M. Comparison of Different Response Time Outlier Exclusion Methods: A Simulation Study. Frontiers in Psychology. 2021;12.

19.	Jakobsen JC, Gluud C, Wetterslev J, Winkel P. When and how should multiple imputation be used for handling missing data in randomised clinical trials – a practical guide with flowcharts. BMC Medical Research Methodology. 2017;17(1).

20.	Thomas Yeo BT, Krienen FM, Sepulcre J, Sabuncu MR, Lashkari D, Hollinshead M, et al. The organization of the human cerebral cortex estimated by intrinsic functional connectivity. Journal of Neurophysiology. 2011;106(3):1125-65.

21.	Hawrylycz MJ, Lein ES, Guillozet-Bongaarts AL, Shen EH, Ng L, Miller JA, et al. An anatomically comprehensive atlas of the adult human brain transcriptome. Nature. 2012;489(7416):391-9.

22.	Markello RD, Arnatkevičiūtė A, Poline J-B, Fulcher BD, Fornito A, Misic B. 2021.

23.	Huang C-C, Rolls ET, Hsu C-CH, Feng J, Lin C-P. Extensive Cortical Connectivity of the Human Hippocampal Memory System: Beyond the “What” and “Where” Dual Stream Model. Cerebral Cortex. 2021;31(10):4652-69.

24.	Arnatkevic̆iūtė A, Fulcher BD, Fornito A. A practical guide to linking brain-wide gene expression and neuroimaging data. NeuroImage. 2019;189:353-67.

25.	Quackenbush J. Microarray data normalization and transformation. Nature Genetics. 2002;32(S4):496-501.

26.	Hawrylycz M, Miller JA, Menon V, Feng D, Dolbeare T, Guillozet-Bongaarts AL, et al. Canonical genetic signatures of the adult human brain. Nature Neuroscience. 2015;18(12):1832-44.

27.	Fulcher BD, Little MA, Jones NS. Highly comparative time-series analysis: the empirical structure of time series and their methods. Journal of The Royal Society Interface. 2013;10(83).

28.	Hastie T, Tibshirani R, Friedman J. Model Assessment and Selection. The Elements of Statistical Learning. Springer Series in Statistics2009. p. 219-59.

29.	Uhlén M, Fagerberg L, Hallström BM, Lindskog C, Oksvold P, Mardinoglu A, et al. Tissue-based map of the human proteome. Science. 2015;347(6220).

30.	Eden E, Navon R, Steinfeld I, Lipson D, Yakhini Z. GOrilla: a tool for discovery and visualization of enriched GO terms in ranked gene lists. BMC Bioinformatics. 2009;10(1).

31.	Hansen JY, Shafiei G, Markello RD, Smart K, Cox SML, Nørgaard M, et al. Mapping neurotransmitter systems to the structural and functional organization of the human neocortex. Nature Neuroscience. 2022;25(11):1569-81.

32.	Gramfort A, Luessi M, Larson E, Engemann DA, Strohmeier D, Brodbeck C, et al. MNE software for processing MEG and EEG data. NeuroImage. 2014;86:446-60.

33.	Widmann A, Schröger E, Maess B. Digital filter design for electrophysiological data – a practical approach. Journal of Neuroscience Methods. 2015;250:34-46.

34.	Bigdely-Shamlo N, Mullen T, Kothe C, Su K-M, Robbins KA. The PREP pipeline: standardized preprocessing for large-scale EEG analysis. Frontiers in Neuroinformatics. 2015;9.

35.	Delorme A, Makeig S. EEGLAB: an open source toolbox for analysis of single-trial EEG dynamics including independent component analysis. Journal of Neuroscience Methods. 2004;134(1):9-21.

36.	Winkler I, Haufe S, Tangermann M. Automatic Classification of Artifactual ICA-Components for Artifact Removal in EEG Signals. Behavioral and Brain Functions. 2011;7(1).

37.	Welch P. The use of fast Fourier transform for the estimation of power spectra: A method based on time averaging over short, modified periodograms. IEEE Transactions on Audio and Electroacoustics. 1967;15(2):70-3.

38.	Donoghue T, Haller M, Peterson EJ, Varma P, Sebastian P, Gao R, et al. Parameterizing neural power spectra into periodic and aperiodic components. Nature Neuroscience. 2020;23(12):1655-65.
